# Conservation of separase N-terminal domain

**DOI:** 10.1101/236216

**Authors:** Michael Melesse, Joshua N. Bembenek, Igor B. Zhulin

## Abstract

We report a reanalysis of the sequence conservation of the cell cycle regulatory protease, separase. The sequence and structural conservation of the protease domain has long been recognized. Here we reexamine the protein sequence conservation at the N-terminus using PSI-BLAST analysis and report our discovery of a cysteine rich motif (CxCXXC) conserved in nematodes and vertebrates. This motif is found in a solvent exposed linker region connecting two TPR-like helical motifs. Mutation of this motif in *Caenorhabditis elegans* separase leads to a temperature sensitive hypomorphic protein, and several N-terminal residues identified as intragenic suppressors are not conserved. Conservation of this motif in multiple organisms raises the possibility that the motif plays similar roles across species.

## Introduction

Separase is a CD clan cysteine protease which cleaves multiple substrates to regulate cell division. Separase proteolytic activity is tightly regulated mainly by the binding of an inhibitory chaperone called securin [1, 2], which is degraded at the onset of anaphase in a proteasome dependent manner after polyubiquitination by the Anaphase Promoting Complex/Cyclosome (APC/C) [3, 4]. Once activated, separase cleaves a subunit of cohesin which holds sister chromatids together, allowing sister chromatids to segregate to opposite poles [5–8]. Later during anaphase, separase cleaves a number of other substrates to regulate the spindle [9], centrosome duplication [10], and exocytosis [11]. Separase is also found to be functionally conserved from single-celled yeast to multicellular mammals with molecular mass ranging from 144 kDa (in *Caenorhabditis elegans*) to 244 kDa (in a *A. thaliana*) [12]. This highly conserved protease is central to the regulation of anaphase in many systems.

Previous reports attempting to define sequence conservation of separase have mainly identified conservation within the C-terminal protease domain of the protein [13–14]. More recent structural analyses of *C. elegans* and yeast separases have found that the geometry of the catalytic active site is also conserved [15–17]. However, separase is a large protein with a sizable N-terminal domain that has been poorly characterized and N-termini of yeast and *C. elegans* are structurally different [16, 17]. *C. elegans* separase is composed of an N-terminal α-solenoid domain connected to the C-terminal caspase-like protease domain [15] (Figure 1). The α-solenoid domain comprises 25 α-helices, mainly arranged as a right-handed superhelix that resembles a TPR superhelix [17]. In contrast to a canonical TPR superhelix, however, the irregular length of its constituent α-helices creates a compact globular structure that lacks the deep surface grooves typical of TPR proteins [17]. Modifications of the N-terminus, including phosphorylation by Cdk1 [18,19], a change in conformation mediated by the peptidyl-prolyl cis/trans isomerase (PPIase) pin1 [20] as well as separase auto cleavage [21, 22] regulate separase activity. Therefore, it is important to understand how the N-terminal domain of separase functions to better understand these different regulatory mechanisms.

**Figure 1:**
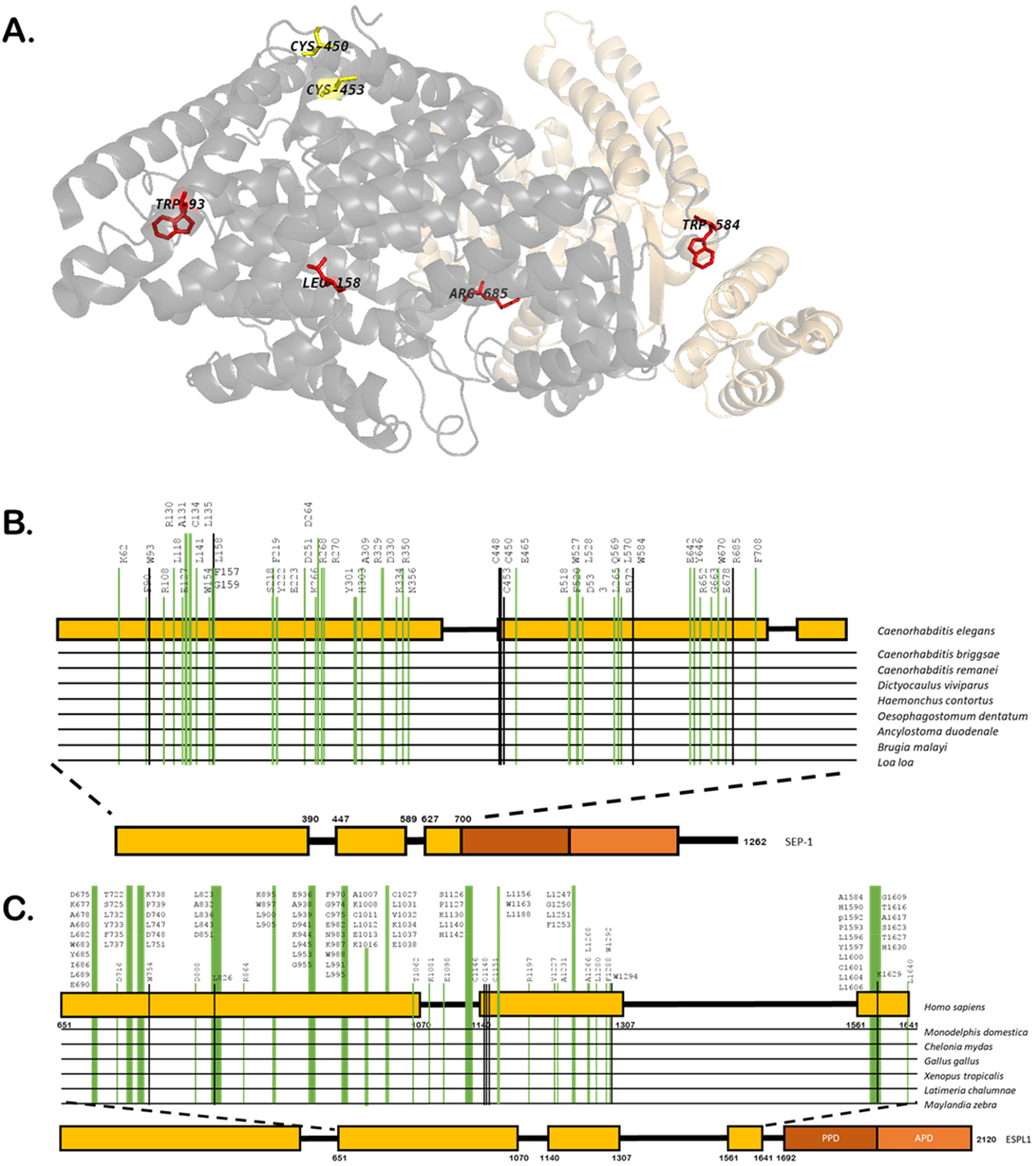
Separase N-terminal domain is conserved across multiple residues. **A**. *C. elegans* separase Cryo-EM structure (PDB 5MZ6) illustrating N-terminal residues universally conserved in nematodes and vertebrates. The N-terminal domain is depicted in gray and C-terminal protease domain is shown in light orange. The two visible conserved residues that make up the cysteine motif (C450 and C453) are shown in yellow and are found as part of a solvent exposed loop between helices 15 and 16, while C448 was not resolved in the crystal structure. Other conserved residues shown in red are distributed across the N-terminus. W93, L158 and R685 are found in the interior of the protein where as W584 is on the protein surface. Different sets of N-terminal residues are conserved among nematode and vertebrate separase. Protein diagram shows N-terminal residues conserved among nematode (**B**) and vertebrate (**C**) separases in green. Residues conserved across all species are shown in black. The residue numbering at the top of each diagram corresponds to *C. elegans* (**B**) and human (**C**) separase.

In *C. elegans*, separase localizes to vesicles and regulates exocytosis events during anaphase [11, 23]. A mutation (Cys450Tyr) in the N-terminus of the *C. elegans* separase (*sep-1(e2406)*) results in a temperature sensitive phenotype that leads to the failure of cortical granule exocytosis (CGE) during anaphase, with little effect on chromosome segregation [11]. During anaphase, the *SEP-1(e2406)* protein localizes to the meiotic spindle, but has reduced localization to cortical granules and supports a lower number of exocytic events. In an extensive suppressor screen, we identified multiple intragenic suppressors of *sep-1(e2406)* that exclusively introduce mutations to the N-terminus [24]. Therefore, we investigated structural conservation in the N-terminus to better understand the structural elements of this domain that are affected by these mutations.

Previous computational studies of separase from different taxonomic lineages revealed no sequence conservation in the N-terminal region corresponding to the α-solenoid domain, whereas the C-terminal region including the caspase-like domain (recognized by the Peptidase_C50 Pfam model) was found to be well conserved [13, 25]. No multiple sequence alignments of the separase N-terminal region are available in the current literature due to the reported extreme divergence; however, a pairwise sequence alignment between the human and nematode separase sequences was produced based on the recently solved three-dimensional structure of the *C. elegans* separase [17]. Twenty-five structurally identified helices, comprising eleven TPR-like repeats in the N-terminal region of the *C. elegans* separase (accession NP_491160.1, aa 1-700), were aligned to the twenty-five α-helices predicted in the N-terminal region of the human separase (NP_036423.4, aa 651-1641). This analysis revealed more than 80 identical residues, including Cys450. The very low percentage of identity between the two sequences and the fact that they were aligned manually, guided by structural information, prompted us to explore potential sequence conservation in the N-terminal region of separase using more conventional bioinformatics approaches.

First, we performed a BLAST [26] search (with default parameters) of the non-redundant protein sequence database at the NCBI (NR) with the N-terminal (aa 1-740) portion of the *C. elegans* separase (accession NP_491160.1) as a query. The search confidently retrieved similar proteins from several distantly related nematode species and reciprocal BLAST searches with full-length sequences validated their orthology. Next, we performed exhaustive PSI-BLAST [27] searches (with default parameters) of the NR database with N-terminal regions of the nematode sequences as queries. The search initiated with the N-terminal region (aa 1-760) of the *Toxicara canis* separase (accession KHN86283.1) retrieved a separase from a vertebrate *Danio rerio* (XP_001337869.1) with E value 9e-04 in the third iteration and a human separase (accession NP_036423.4) with E value 2e-11 in the fourth iteration, among many other separase sequences from vertebrates. While having statistically significant matches to the PSI-BLAST generated profile, vertebrate separase sequences showed low identity (10-12%) to nematode sequences. We then constructed a multiple sequence alignment of representative nematode and vertebrate sequences using MAFFT [28] with default parameters (Supplementary Figure 1). Overall, the alignment, which was only minimally edited based on structural information, looks similar to that of the *C. elegans* and *H. sapiens* sequences published by [17]; however, there are drastic differences with respect to conserved positions revealed by these alignments. While more than 80 identical residues in the *C. elegans* and *H. sapiens* sequences were identified by (Supplementary Figure 4 of [17]), our alignment shows only 7 positions that are identical in all nematode and vertebrate sequences. The majority of identical positions revealed by comparing only two sequences do not match conservation patterns revealed by our multiple sequence alignment. These alignments appear to be due to incorrect gap placements or happen purely by chance (Supplementary Figure 1), which further emphasizes problems associated with building and interpreting “manual alignments”, even when assisted by structural information [29]. Among the very few identical positions that we identified in nematode and vertebrate N-terminal sequences (Figure 1A), four are in various parts of the region: W93/W754 (*C. elegans/H. sapiens*), L158/L827, W584/W1294, and R685/R1629. The remaining three identical residues, C448/C1146, C450/C1148, and C453/C1151, appear to form a motif, which is located at the border of Insert 1 and helix H16 (see Supplementary Figure 1 for details). Other conspicuous conserved positions include: F90/L751, E127/K789, F219/W898, K262/D940, D330/E1006, and F520/Y1227. As expected, multiple residues, distributed throughout the N-terminal domain (Supplementary Figure 2), are conserved among nematode separases; which are different from residues conserved among vertebrate separase (Figure 1B and 1C). None of these conserved residues are in contact with the separase inhibitory chaperone, securin. The nature of this conservation remains unclear. None of the highly conserved residues appear to form a contact with another similarly conserved residue, as judged by the proximity of beta carbons on the separase structure (Supplementary Table 1). However, their functional importance is highlighted by the fact that no changes in these positions are observed in human populations (Supplementary Figure 3) and mutation in one of them, C450, results in temperature sensitive embryonic lethality in *C. elegans* [11, 23]. Residues that can, when mutated, suppress *sep-1(e2406)* [24] are not conserved, do not contact securin and are distributed throughout the N-terminus (Supplementary Figure 2C). Understanding effects of these mutations on SEP-1 structure will require further investigation. Our analysis demonstrates the utility of studies in *C. elegans* in understanding separase regulation in multiple organisms.

---

**Supplementary Figure 1.**
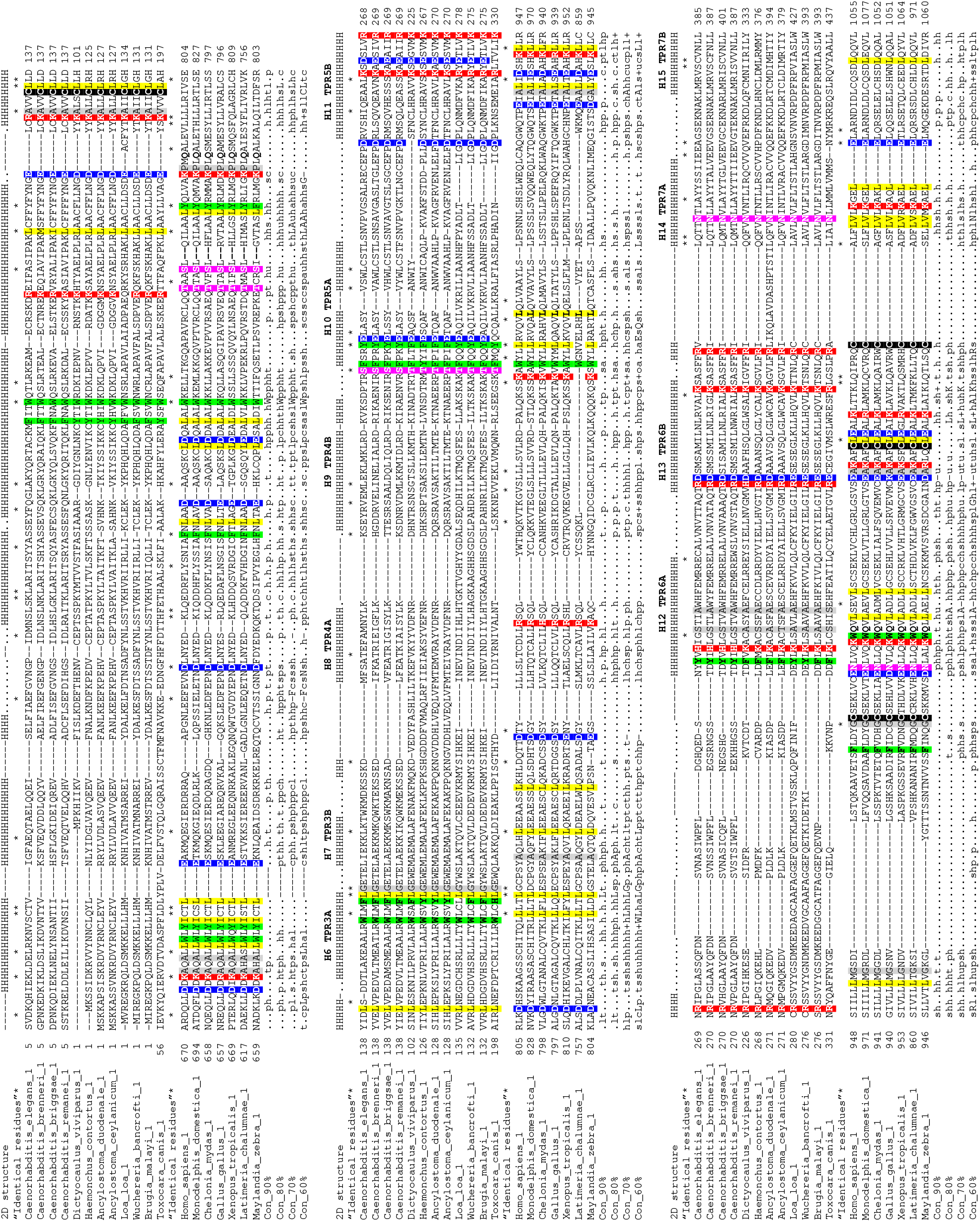
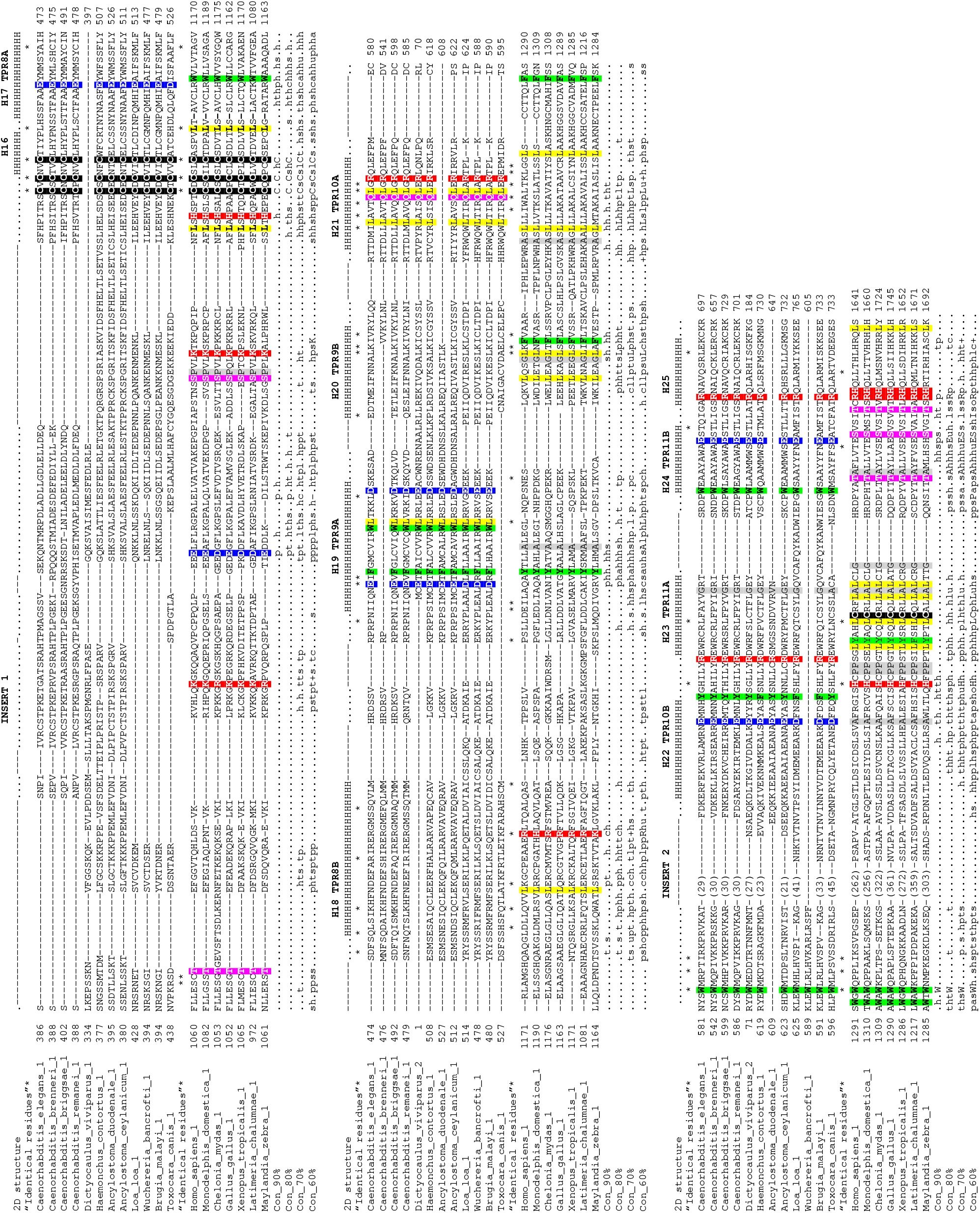
Multiple sequence alignment of separases from nematodes and vertebrates. Sequences from eleven nematode species (top portion of each panel) and from seven representative vertebrate species (human, opossum, turtle, chicken, frog, coelacanth, and fish) are shown. Twenty-five alpha helices (labeled H1 to H25) comprising eleven TPR-like repeats (labeled TPR1A,B to TPR11A,B) in the *C. elegans* separase (PDB accession 5MZ6) are shown above the alignment. Identical residues in each group are highlighted: negatively charged, red; positively charged, blue; aromatic, green; aliphatic, yellow; alcohol, magenta; small, grey. NCBI accession numbers: Caenorhabditis_elegans_1, NP_491160.1; Caenorhabditis_brenneri_1, EGT38506; Caenorhabditis_briggsae_1, CAP33358; Caenorhabditis_remanei_1, XP_003114963.1; Loa_loa_1, XP_003140515.1; Wuchereria_bancrofti_1, EJW80934; Brugia_malayi_1, XP_001894870.1; Dictyocaulus_viviparus_1, KJH53363.1; Dictyocaulus_viviparus_2, KJH53362.1; Haemonchus_contortus_1, CDJ83415.1; Ancylostoma_duodenale_1, KIH65515.1; Ancylostoma_ceylanicum_1, EYC45610.1; Toxocara_canis_1, KHN86283.1; Homo_sapiens_1, NP_036423.4; Monodelphis_domestica_1, XP_007506592.1; Chelonia_mydas_1, XP_007058605.1; Gallus_gallus_1, XP_015128534.1; Xenopus_tropicalis_1, XP_004912005.1; Latimeria_chalumnae_1, XP_014347491.1; Maylandia_zebra_1, XP_014264400.1. “Identical residues”* show positions defined as identical in a pairwise comparison of *C. elegans* and *H. sapiens* sequences by Boland et al., 2017.

**Supplementary Figure 2:**
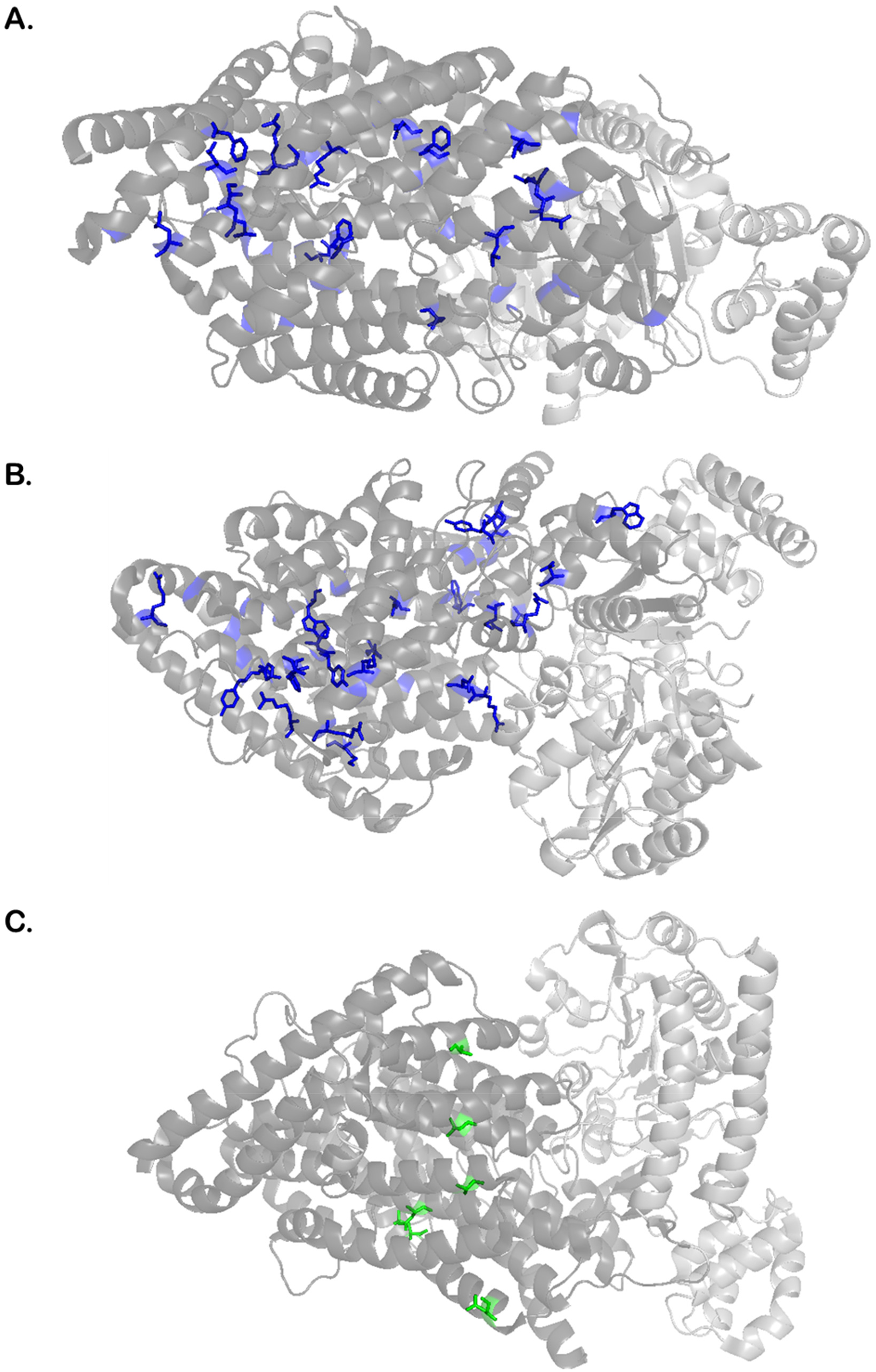
Separase N-terminal residues conserved among nematodes are distributed throughout the structure. *C. elegans* separase Cryo-EM structure (PDB 5MZ6) illustrating N-terminal residues conserved among nematodes found in the interior (**A**) and on the surface (**B**) of the TPR-like N-terminal domain. Intragenic suppressors of SEP-1(e2406) are shown (**C**) and are not among the conserved residues. The structures are oriented with the N-terminus to the left with a perspective that best illustrates the distribution of each highlighted residue.

**Supplementary Figure 3:**
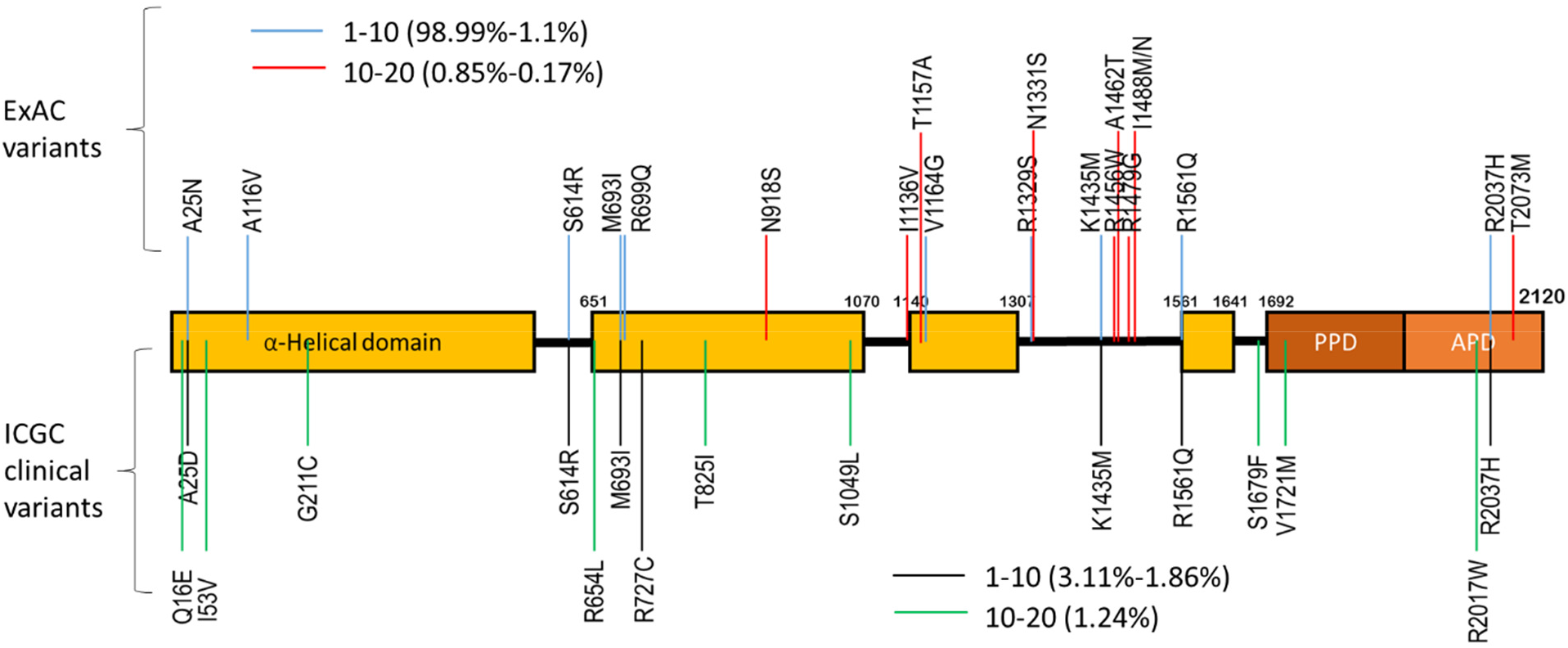
Mutations in human separase (ESPL-1) associated with cancers. The collection of separase allelic variants of human Separase from the ExAC collection (http://exac.broadinstitute.org) and the ICGC (https://icgc.org/) which collects genomic sequences of various cancers. The frequency of each missense mutation are indicated.

**Supplementary Table 1:** Conserved residues found within helices that makeup the N-terminal domain are listed. Residues that are within interacting distance (as assayed by measuring a distance less than 6 angstroms (Å) between β-Carbons) are indicated. These residues are generally located within the same helix and don’t appear to be important for stabilizing inter-helix interactions.

| Conserved residue | <6 Å β-C distance | Same helix? |
| --- | --- | --- |
| F90 | W93 (5.5 Å) | same |
| W93 | F90 (5.5 Å) | same |
| E127 | R130 (4.7 Å) | same |
| L158 | I514 (5.3 Å) | no |
| F220 | Y222 (6.0 Å) | same |
| D264 | None |  |
| D330 | R329 (5.1 Å) | same |
| F520 | None |  |
| W584 | Y582 (5.7 Å) | same |
| R685 | None |  |

